# The camtrapR R package: From data management to interactive ecological analysis of camera trap data

**DOI:** 10.1101/2025.09.26.678697

**Authors:** Jürgen Niedballa, Rahel Sollmann, Andreas Wilting

## Abstract

1. Camera trapping has become an indispensable tool in wildlife ecology, generating vast datasets that require efficient and robust analytical workflows. The R package `camtrapR` was originally developed for preparing and managing camera trap data for subsequent analysis in external modeling packages like `unmarked`. It has since become a standard tool in the field for this purpose.
2. Here, we introduce a major update that transforms `camtrapR` from a data preparation tool into a comprehensive, end-to-end analytical platform. The centerpiece of this evolution is the surveyDashboard(), a novel code-free graphical user interface that guides users through the entire analysis pipeline, from data import to final predictions. This update also incorporates enhanced data import functionalities for major standards like Wildlife Insights and Camtrap DP, a complete workflow for fitting community occupancy models, and streamlined tools for environmental covariate extraction.
3. The interactive dashboard provides an integrated environment for the entire analytical process. Users can perform essential exploratory analyses, such as generating species accumulation curves and mapping species detections, before proceeding to model fitting. The interface supports the interactive construction of both single-species and multi-species (community) occupancy models. The dashboard’s covariate preparation tools generate inputs for both model fitting and for creating spatial predictions of species occupancy.
4. Furthermore, the update introduces a comprehensive workflow for fitting Bayesian community occupancy models using JAGS or NIMBLE. This allows for hierarchical modeling of species- and community-level responses to environmental drivers, providing deeper insights into wildlife communities. The workflow includes tools for model assessment, such as convergence diagnostics and posterior predictive checks for goodness-of-fit.
5. By integrating a powerful, code-free interface with advanced backend modeling functions, this major update to `camtrapR` aims to make robust and reproducible camera trap data analysis accessible to a wider audience, including ecologists, wildlife managers, and students. This paper serves as the new definitive reference for the expanded functionality of `camtrapR` as a comprehensive tool for modern camera trap studies.

## Introduction

Camera trapping has transformed wildlife research and has become an indispensable tool for studying terrestrial wildlife (Wearn and Glover-Kapfer 2019). The method now supports diverse applications, from species inventories to behavioral studies and from individual species distribution studies to community assessments (Delisle et al. 2021). Yet with widespread adoption has come the challenge of managing and analyzing the substantial volumes of data these devices generate (Steenweg et al. 2017).

We first released the R package `camtrapR` on CRAN in 2015 to address the growing need for standardized data management in camera trap studies (Niedballa et al. 2016). The R package streamlined data organization and preparation for occupancy and spatial capture-recapture analyses. It found broad adoption among researchers, particularly those requiring reproducible data management workflows (Thomson et al. 2018). However, the field has evolved considerably, creating new demands that extend beyond the original package’s capabilities.

Several developments in camera trap research now require enhanced analytical tools. The emergence of standardized platforms like Wildlife Insights and Camera Trap Data Package (camtrap DP) has created new data-sharing frameworks that need support (Bubnicki et al. 2024; Ahumada et al. 2020). There is also increasing focus on community-level analyses, reflecting the need to understand species interactions and community-level responses to environmental variables (Devarajan et al. 2020; Tingley et al. 2020). Additionally, the integration of spatial data in species distribution modeling has become standard practice, requiring more sophisticated tools for handling environmental covariates.

To date, no platform integrates the full workflow from camera trap data management to sophisticated ecological analyses. Existing solutions often address only portions of this workflow: Wildlife Insights and TrapTagger function primarily as a data repositories with AI-based species identification while developing analytical capabilities; platforms like Agouti, Camelot and TRAPPER (Enetwild Consortium et al. 2022; Hendry and Mann 2018; Bubnicki et al. 2016) offer data management with basic analyses; specialized software such as AddaxAI and Sherlock (Penn et al. 2024) focus on AI-based species identification; Camera Trap Data Package (camtrap DP, Bubnicki et al. 2024) provides a metadata standard without analytical tools; and statistical R packages like `unmarked`, `ubms` and `spOccupancy` (Kellner et al. 2023; Kellner et al. 2022; Doser et al. 2022) offer powerful analytical capabilities for single and multi-species models, but lack camera trapping-specific data handling. The updated `camtrapR` package addresses this fragmentation by offering a complete workflow that connects camera trap data management tools with analytical packages while introducing new features not available elsewhere, particularly community occupancy models that allow simultaneous analysis of multiple species while accounting for imperfect detection in a graphical user interface.

The user base for camera trap data has also expanded beyond academia to include conservation practitioners, wildlife managers, and citizen scientists (Willi et al. 2019). This diversification creates a need for more accessible tools that maintain analytical rigor while reducing technical barriers.

In this paper, we present significant updates to `camtrapR` that address these evolving needs. The package now features an interactive Shiny dashboard for code-free analysis, enhanced data import capabilities, integrated community occupancy modeling, and streamlined handling of spatial covariates, among other improvements. These developments maintain the package’s commitment to reproducible workflows while making advanced analytical methods accessible to a broader user base. We demonstrate these new capabilities through a worked example and discuss their implications for future camera trap research.

## New Functionalities and Improvements

The updated `camtrapR` package introduces key enhancements that expand its analytical capabilities while improving accessibility. We focused developments in three main areas: interactive analysis, expanded analytical tools, and enhanced data handling.

### Interactive Analysis Dashboard

The most significant addition is an interactive dashboard that allows users to analyze camera trap data without writing code. Built using R’s Shiny framework (Chang et al. 2025), the dashboard integrates data import, exploration, and analysis in a unified interface. It contains help pages for all major components and tooltips for all UI elements to guide users, and provides immediate visual feedback as they explore their data, generate summaries, and build statistical models. We first provide an overview of the dashboard and below more detailed descriptions of the novel features, which are also available via standalone functions outside the dashboard.

The dashboard comprises several interconnected modules. Users can import their data from various sources (CSV files, WildlifeInsights, camtrap DP). The summary tab shows key information about surveys and includes interactive maps of the camera trapping grid. Users can filter their data dynamically by site covariates, or filter records by species or temporal independence. Users can visualize species activity patterns and species accumulation curves (Meredith and Ridout 2016; Hsieh et al. 2016). A dedicated covariate extraction module helps users create covariates from spatial rasters and from online elevation databases for use in distribution modeling. It supports the creation of derived terrain metrics and harmonizes all raster data for spatial predictions (Hijmans et al. 2025). Users can further examine relationships and correlation between environmental variables.

The modeling interface guides users through building occupancy models, supporting both frequentist and Bayesian approaches through the `unmarked` and `ubms` packages for single-species analyses. Additionally, the dashboard includes a dedicated interface for building and fitting community (multi-species) occupancy models, allowing users to analyze multiple species simultaneously while accounting for imperfect detection. Advanced features of the occupancy workflows include model comparison, visualization of species responses to environmental variables, and creation of spatial predictions.

### Enhanced Data Import and Standardization

To support emerging data standards, we added dedicated import functions for Wildlife Insights and Camera Trap Data Package (camtrap DP) formats to both the package and the dashboard (Ahumada et al. 2020; Bubnicki et al. 2024). These functions handle the specific structure and metadata requirements of these platforms automatically.

Besides camera trap photos, the package also supports video data, though extracting metadata from videos remains challenging due to inconsistent standards across camera brands.

### Expanded Analytical Capabilities

#### Species accumulation curves

To assess sampling completeness, `camtrapR` integrates the rarefaction and extrapolation framework of the `iNEXT` package (Chao et al. 2014; Hsieh et al. 2016). We extend the standard `iNEXT` application by enabling analysis along both spatial and temporal dimensions. The spatial approach, a standard application of `iNEXT`, uses camera trap locations as sampling units to evaluate how species diversity increases with sampling locations. Complementing this, `camtrapR` introduces a novel temporal analysis where survey days (or camera-specific operational days) are treated as sampling units. While the spatial analysis assesses the sufficiency of site coverage, our novel temporal implementation allows researchers to evaluate sampling completeness as a function of survey duration. This dual functionality provides a more robust assessment of survey effort, with confidence intervals to quantify uncertainty in sampling completeness and inform study design.

#### Automated Spatial Covariate Handling

The new `createCovariates` function simplifies the integration of spatial data into analyses. It extracts covariate values from local raster files or online elevation databases, manages coordinate system transformations, and prepares standardized inputs for both model fitting and prediction. This automation drastically reduces the technical expertise needed to incorporate environmental data while ensuring consistent spatial data handling. Elevation data are obtained from Amazon Web Services (AWS) Terrain Tiles via the `elevatr` package (Hollister et al. 2023). For a selection of data sources for use as environmental covariates, see Supporting Table S1.

#### Community Occupancy Models

A major analytical addition is support for community occupancy models through the `communityModel` function. This implementation handles both standard occupancy models and Royle-Nichols abundance-based occupancy models, allowing users to specify various covariate effects on detection and occupancy probabilities. Models can be fitted using either JAGS or Nimble, with specialized functions for examining results (`summary`), visualizing effects (`plot_effects`), and making spatial predictions (`predict`). The framework incorporates random effects of species and sites, enabling analysis of community responses to environmental conditions while accounting for species-specific variations.

#### Additional Improvements

We added several other enhancements to improve the package’s utility. For example, the `filterRecordTable` function provides flexible temporal filtering of detection records, useful when importing external data. Camera operation calculations now account for precise timing of camera deployment and retrieval. For situations where image metadata is unavailable, new optical character recognition capabilities can extract date and time information directly from images (Smith 2007; Ooms 2025).

## Worked Example

To demonstrate `camtrapR`’s new capabilities, we analyze a subdataset of the SNAPSHOT USA data set (Rooney et al. 2025) collected in Lubrecht Experimental Forest, Montana (USA), where 31 locations were surveyed with 57 camera deployments from September to December 2022 to study the effect of hunting on ungulates and pumas. The dataset includes 1602 independent detection events of 24 identifiable wildlife species.

Our objective here is to examine how species occurrence patterns and community composition vary with elevation and terrain characteristics across the landscape. The manuscript presents a high-level overview, and Supporting Information S2 is an R markdown document containing a fully worked, executable example.

### Alternative Workflows

Camera trap data can be analyzed through two complementary approaches in `camtrapR`. The interactive dashboard supports data exploration, visualization, interactive model development and model predictions. It is particularly valuable during initial analysis phases, yet capable of sophisticated analyses. The code-based workflow facilitates reproducibility and automation, suitable for standardized, reproducible analyses or processing multiple datasets in a consistent way. We demonstrate how to load data, add site covariates from online sources and analyse species habitat associations with a multi-species occupancy model using both approaches to highlight their utility in different contexts. Figure 1 gives an overview of the workflow and example outputs.

**Figure 1.**
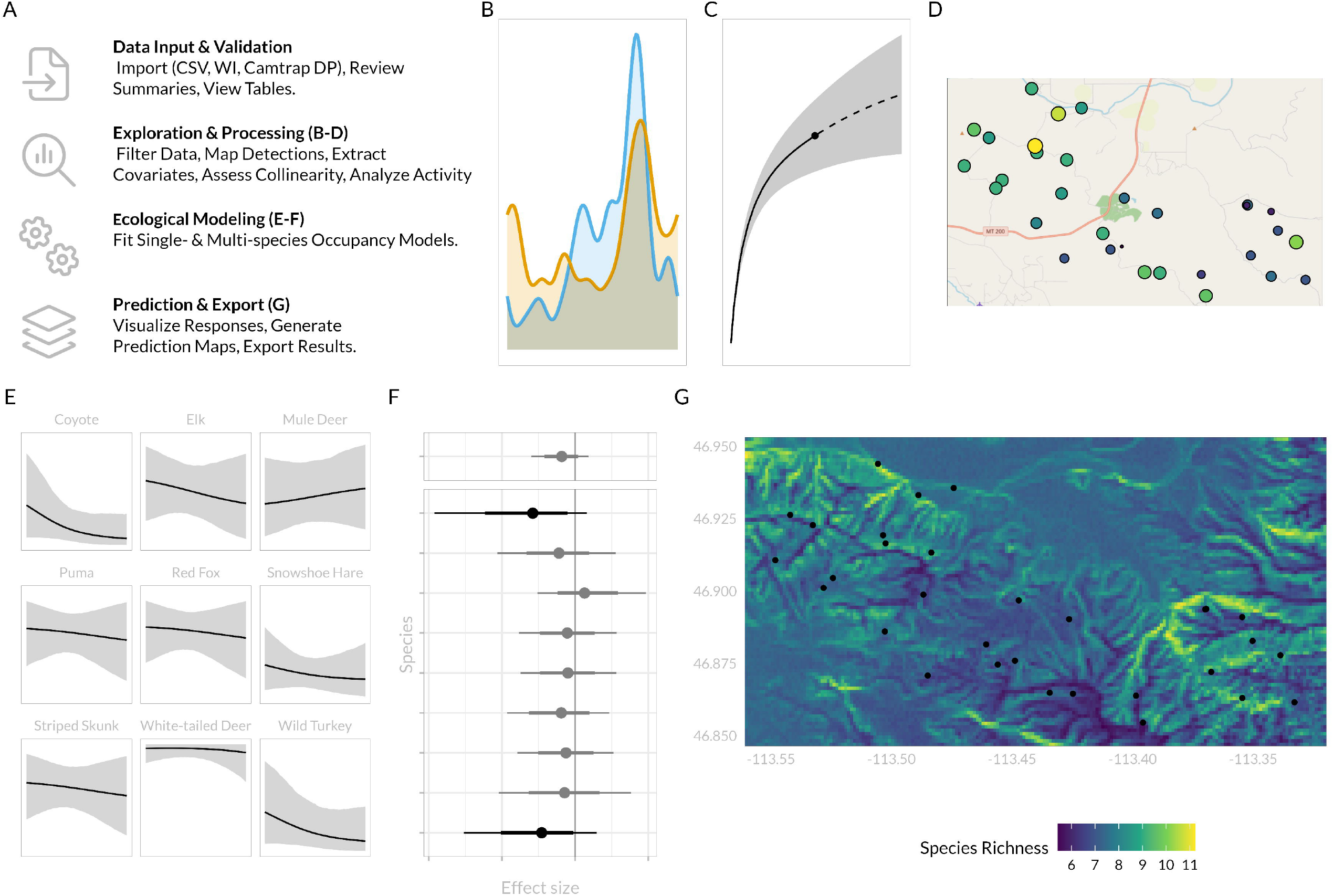
A conceptual workflow and example outputs from the camtrapR interactive dashboard. (A) The dashboard guides users through four main stages of analysis, from data import to prediction. Key modules are grouped within each stage. Example exploratory outputs include (B) diel activity pattern overlap plots, (C) species accumulation curves and (D) maps of observed species richness. (E) The modeling interface allows users to fit single- and multi-species occupancy models and visualize model results, such as the relationship between occupancy probability and an environmental covariate and (F) the effect size by species. (G) Final model outputs can be used to generate spatial predictions, such as this map of predicted species richness for a species community (with camera locations overlaid).

### Dashboard-Based Analysis

After launching the dashboard with

`surveyDashboard()`, or pre-loading it with specific parameters (to avoid the initial data import), the analysis proceeds through several steps, all of which are implemented on the dashboard interface and accessible without any coding.

#### 1. Data Import and Validation

- **Import Raw Data: Load** camera trap data from various formats (CSV, Wildlife Insights, Camtrap DP) using the “Import” module, or by passing parameters directly to surveyDashboard().
- **Review and Validate: Review** summary statistics, sampling effort, and camera operational timelines in the “Data Summary” module to confirm data integrity. **Verify** correct spatial referencing by examining station maps in the “Maps” module.
- **Prepare Covariates: Extract** environmental covariates by combining automatically derived terrain metrics with user-provided local raster files (see Supporting Table S1). **Assess** collinearity using correlation plots and **select** a final set of non-collinear covariates for modeling

#### 2. Exploratory Data Analysis and Filtering

- **Visualize Ecological Patterns: Generate** species accumulation curves to assess sampling adequacy. **Map** species detections and observed richness to visualize spatial patterns (“Maps” module). **Visualize** species activity patterns to explore diel activity and temporal overlap (“Species Activity” module).
- **Filter and Subset: Filter** the dataset by stations, species, or covariate values using the “Data Filters” module to create subsets for specific analyses.

#### 3. Occupancy Modeling

- **Define and Fit Models: Select** between single-species (unmarked/ubms) and multi-species (JAGS/NIMBLE) occupancy models. **Define** detection histories and **specify** the model structure by interactively selecting covariates (and random effects for multi-species models) for detection and occupancy parameters. **Run** the model fitting process.
- **Assess Model Performance: Examine** MCMC convergence diagnostics and **run** posterior predictive checks for goodness-of-fit. **Build** and compare models using a model selection table.

#### 4. Prediction, Visualization and Export

- **Visualize Model Results: Plot** community- and species-level responses to covariates (marginal effect plots).
- **Generate Predictions: Create** prediction maps for individual species occupancy and community-level species richness. **Estimate** derived parameters, such as the percentage of area occupied (PAO).
- **Export: Export** all tables, figures, and model objects for reporting and further analysis.

### Code-Based Implementation

All functionalities of the graphical user interface are accessible programmatically, enabling reproducible, script-based analyses with greater flexibility. The package’s functions are designed to create a logical and intuitive pipeline, mirroring the analytical workflow. We illustrate this core pipeline below, noting that each function contains extensive customization options detailed in the package documentation. Full, executable code for this example is provided in the Supporting Information S2.

First, we load data and prepare inputs. The `readWildlifeInsights()` function is one of several ways to import data into `camtrapR` and streamlines data import by automatically processing the zip file exported by WildlifeInsights.

~~~
wi_data <-readWildlifeInsights(“survey_export.zip”)
~~~

Next, `createCovariates()` prepares the environmental data by combining local raster files and online elevation and terrain data.

~~~
covs <-createCovariates(CTtable = wi_data$stations,
           directory = “/path/to/covariates”
           download_elevation = TRUE,
           terrain_measures = c(“slope”, “roughness”))
~~~

Camera operational periods are then summarized with `cameraOperation()`.

~~~
camop <-cameraOperation(CTtable = wi_data$stations)
~~~

And finally, detection histories are compiled using detectionHistory() for selected species and a defined occasion length (in days).

~~~
DetHist_list <-detectionHistory(recordTable = wi_data$records,
                     camOp = camop,
                     species = c(“species_1”, “species_2”),
                     occasionLength = 10)
~~~

Detections, site and observation covariates are then bundled in a single list for use as model input.

~~~
model_input <-list(ylist = DetHist_list,
             siteCovs = covs$CTtable,
             obsCovs = list(effort = DetHist_list$effort))
~~~

With the data prepared, users can define and fit various community occupancy models. The `communityModel` function integrates detection histories, site covariates, and observation covariates into a single data structure, allowing for flexible model specification. The model is then fitted using Markov Chain Monte Carlo (MCMC) simulations.

~~~
comm_mod <-communityModel(data_list = model_input,
                 occuCovs = list(ranef = c(“elevation”, “slope”)),
                 detCovsObservation = list(fixed = “effort”))
fit <-fit(comm_mod, n.iter = 10000, chains = 3)
~~~

Predictions can be generated from the model output and environmental covariates.

~~~
pred_richness <-predict(object = comm_mod,
              mcmc.list = fit,
              type = “richness”,
              x = covs$predictionRaster)
~~~

The `predict` function’s output format is determined by the `type` argument. It returns spatial rasters for predictions of species richness, occupancy probability, or abundance (from Royle-Nichols models) containing layers for the mean, standard deviation, and confidence intervals. Alternatively, it can also calculate Percentage of Area Occupied (PAO), a summary statistic which is returned as a data frame.

### Key Results

As a demonstration of `camtrapR`’s modeling capabilities, we fitted a community occupancy model to the example Snapshot USA dataset. While all model steps, from data preparation to prediction, were successfully executed, the 95% credible intervals of species’ responses to environmental covariates like elevation, ruggedness, and topographic position did not show evidence for strong positive of negative effects in this particular small dataset and for the presented model structure. Some species showed trends (75% credible intervals not overlapping zero) and these are presented to illustrate the output format and interpretation of coefficient plots generated by the package.

This example demonstrates how `camtrapR`’s new features integrate to provide a comprehensive analytical workflow, with the option of a visual and interactive dashboard offering robust ecological inference while maintaining accessibility, and traditional, reproducible code-based analysis for users requiring maximum control and reproducibility. Both workflows deliver identical results.

## Discussion

The enhanced capabilities of `camtrapR` reflect the evolving needs of the camera trap research community. By combining accessible interactive tools with sophisticated analytical methods, the package helps bridge the gap between data collection and robust ecological inference. The integration of standardized data formats, community-level analyses, and spatial data handling addresses key challenges in contemporary camera trap research.

The new features respond to distinct needs in the research community. The interactive dashboard lowers technical barriers to statistical analyses while encouraging exploration of ecological patterns. This is particularly valuable during initial data assessment and for users new to occupancy modeling, for whom it can also serve as a learning and teaching tool. The community modeling framework allows researchers to leverage data from all detected species while accounting for imperfect detection, and facilitating access to advanced multi-species occupancy models. Automated handling of spatial covariates considerably simplifies the integration of environmental data into analyses without the need for specialist GIS skills.

Beyond its research applications, the package serves as an effective teaching tool. The immediate visual feedback provided by the dashboard helps users develop intuition about occupancy modeling and species-environment relationships. This aspect is particularly valuable for building analytical capacity in regions where camera trapping is increasingly used for biodiversity monitoring, but statistical expertise may be limited (Stephenson 2020).

While the dashboard makes complex analyses more accessible and provides basic guidance, we emphasize that users should understand the underlying statistical concepts in occupancy modeling before interpreting results for scientific publications or management decisions. We recommend that users unfamiliar with these methods consult statistical experts or relevant literature to ensure appropriate model specification and interpretation.

The current implementation has some limitations that suggest future development priorities. Community occupancy models can be computationally demanding for datasets with many species and sites, or when predicting over large areas and/or at high resolution. Support for video files remains limited by inconsistent metadata standards across camera brands. While `camtrapR`’s outputs are well-suited for standard single-season occupancy and Royle-Nichols models, a key future direction is to unlock the full analytical potential offered by specialized modeling packages. We will develop dedicated coercion functions to provide seamless data handoffs to `unmarked`, `ubms`, and `spOccupancy`. This will enable users to easily apply a more diverse suite of advanced models—including multi-season dynamic, multi-species, and spatial occupancy frameworks—directly to their `camtrapR`-prepared data. Another potential avenue is the inclusion of or interfacing with machine learning tools to help with species identification, which remains one of the most time-consuming steps in camera trap data management and which is an area of intense development (see e.g. Vélez et al. 2023; Beery et al. 2019; Norouzzadeh et al. 2017; Leorna and Brinkman 2022). By providing import functionality for data that were processed by machine learning tools, `camtrapR` makes these data available for analysis in R.

We encourage users to contribute to `camtrapR`’s development through the package’s GitHub repository. User feedback has shaped the current features and will guide future improvements. As camera trap technology and analytical methods advance, we remain committed to maintaining and enhancing `camtrapR`’s utility for wildlife research and monitoring.

## Data availability

The `camtrapR` package is freely available from the Comprehensive R Archive Network (CRAN) at https://CRAN.R-project.org/package=camtrapR. The development version, including the latest features and bug fixes, can be accessed through GitHub (https://github.com/jniedballa/camtrapR). The package includes comprehensive documentation through five detailed vignettes covering image organization, species identification, data extraction, visualization, and multi-species occupancy modeling. Sample datasets and code demonstrating key functionalities are included in the package. The `camtrapR` Google group (https://groups.google.com/g/camtrapr) serves as a help and support forum for users.

All code used in the worked example is provided in the Supporting Information S2. The example dataset used in this paper is available through Dryad at https://doi.org/10.5061/dryad.k0p2ngfhn.

## Acknowledgements

We thank An The Truong Nguyen for extensive testing of the package and providing valuable feedback that substantially improved its functionality. We also appreciate the testing contributions from Ahmed Manzar Baqai, Seth T. Wong, Andrew Tilker, Mwezi Mugerwa, Roshan Guharajan and Yannis Alexiou. This work was supported by the United States Agency of International Development (Biodiversity Conservation project in Viet Nam, 7204402CA00001) and the Bundesministerium für Forschung, Technologie und Raumfahrt (SmartPatrol project, 16LW0439).

