## Supporting Table S1 for "The camtrapR R package: From data management to interactive ecological analysis of camera trap data"

### Supplementary Table S1: Selected Remote Sensing Data Sources for Ecological Covariates

| **Data Source** | **Data Type** | **Spatial Resolution** | **Temporal Coverage** | **Access Method in camtrapR** | **Key Features** |
| --- | --- | --- | --- | --- | --- |
| AWS Terrain Tiles | Elevation, derived terrain metrics | 30m globally | Static | Direct through createCovariates() function with download_elevation = TRUE and terrain_measures defined | Provides elevation data and derived metrics (slope, aspect, roughness, TPI, TRI) without requiring additional downloads |
| GEE-PICX^1^ | Sentinel-2 and Landsat 5-9 multispectral imagery | 10-30m | 1984-present | Local raster import via createCovariates() after download from GEE-PICX | Cloud-free composites, multiple spectral indices (NDVI, EVI, NDWI, etc.), seasonal and annual aggregation options |
| Hansen Global Forest Change^2^ | Forest cover and loss | 30m | 2000-2024 | Local raster import via createCovariates() | Year of forest loss, percent tree cover |
| WorldPop^3^ | Human population density | 100m | 2000-2020 | Local raster import via createCovariates() | Annual population estimates |
| CHELSA^4^ | Climate data (temperature, precipitation, bioclimatic variables) | ~1km | 1979-present | Local raster import via createCovariates() | High-resolution climate data, future climate projections available |
| SoilGrids^5^ | Soil properties (texture, pH, organic carbon, etc.) | 250m | Static | Local raster import via createCovariates() | Global soil information at multiple depths |
| OpenStreetMap | Road networks, settlements, waterways | Vector data | Current | Convert to raster before using with createCovariates() | Infrastructure and human presence features |
| Least cost paths | Slope-dependent cost of movement across the terrain from point sources | custom | Current (if based on OSM) | Local raster import via createCovariates() after processing with e.g. movecost R package | Provides travel time from start points to the landscape, e.g. for assessment of remoteness (walking time from access points) |

**^1^** [**https://github.com/EcoDynIZW/GEE-PICX**](https://github.com/EcoDynIZW/GEE-PICX)

**^2^** [**https://glad.earthengine.app/view/global-forest-change**](https://glad.earthengine.app/view/global-forest-change)

**^3^** [**https://www.worldpop.org/**](https://www.worldpop.org/)

**^4^** [**https://chelsa-climate.org/**](https://chelsa-climate.org/)

**^5^** [**https://soilgrids.org/**](https://soilgrids.org/)

**Note:** To use local raster files with camtrapR, download the data from these sources and optionally process as needed before providing the directory, file path or file names to the createCovariates() function. The function automatically handles coordinate system transformations to provide consistent output. The dashboard interface in camtrapR supports interactive visualization of covariate rasters and assessment of their correlation (at camera trap locations) before inclusion in analytical models.

Vector data sources require conversion to raster format. For example, users may download road data from OpenStreetMap and use these as input for distance rasters or for least-cost-path analyses (e.g. with the R package movecost) to assess travel time from roads to anywhere in the landscape.
