## Supporting Information S2 for "The camtrapR R package: From data management to interactive ecological analysis of camera trap data": camtrapR publication - Supporting Information S2.html

Supplementary Material: Full Analysis Code for the community occupancy workflow in camtrapR


Code 

- Show All Code
- Hide All Code

### Supplementary Material: Full Analysis Code for the community occupancy workflow in `camtrapR`

###### Jürgen Niedballa

#### 2025-09-16

### Introduction and Setup

This supplementary material provides the complete R code used to
perform the analyses described in the main manuscript, demonstrating the
functionality of the `camtrapR` R package. The code covers
data loading, pre-processing, model definition, fitting, and
prediction.

**To reproduce this analysis:**

1. **Download all files** from the CSV files
   (`ssusa_finaldeployments.csv`,
   `ssusa_finalsequences.csv`) from the SNAPSHOT USA Dryad data
   repository: `https://doi.org/10.5061/dryad.k0p2ngfhn`
2. **Ensure all files are in the same directory.** This R
   Markdown document (
   `supplement_camtrapR_community_analysis.Rmd` ) expects a
   specific directory structure:
   - A `data/` subdirectory containing the csv files.
   - A `figures/` subdirectory containing
     `dashboard_screenshot1.png` (for visual demonstration).
3. **Open the `camtrapR_supplement.Rproj`
   file** in RStudio. This will set the correct working
   directory.
4. **Install necessary packages** (listed below).
5. **Run (knit) this
   `supplement_camtrapR_community_analysis.Rmd`
   file.**

**Important Disclaimer Regarding Ecological
Interpretation:** The data used in this demonstration is derived
from Snapshot USA, and the models are run to showcase the
`camtrapR` package’s capabilities and workflow. While the
underlying model results are derived using sound statistical methods,
this demonstration **does not constitute a full ecological study
or analysis and omits important ecological variables.**
Consequently, **no ecological conclusions, population inferences,
or conservation recommendations should be drawn from these illustrative
outputs.** These results are presented solely to exemplify the
package’s usage.

Please ensure you have the most recent version of
`camtrapR` installed and loaded, along with any other
necessary R packages. For installation instructions please see the
GitHub page (https://github.com/jniedballa/camtrapR).

```
library(camtrapR)
library(sf)
library(terra)
library(unmarked)
library(mapview)     # optional, for interactive plots
library(readr)
library(ggplot2)
library(bayesplot)
library(DT)
```

### Data source

This demonstration uses a small subset of data from SNAPSHOT USA
which is available from Dryad (Rooney, McShea, et
al., 2025). The full SNAPSHOT USA dataset is described in Rooney, Kays, et al. (2025). The subset used in
this demonstration was collected in Lubrecht Experimental Forest,
Montana, USA, by Parsons (2023).

### Load data

The examples uses the two CSV files
(`ssusa_finaldeployments.csv` and
`ssusa_finalsequences.csv`) which can be downloaded from
Dryad.

```
cams_all <- read.csv("data/ssusa_alldeployments.csv")
recs_all <- read.csv("data/ssusa_allsequences.csv")
```

Inspect the data.

```
str(cams_all)
```

```
## 'data.frame':    9694 obs. of  13 variables:
##  $ Year             : int  2019 2019 2019 2019 2019 2019 2019 2019 2019 2019 ...
##  $ Project          : chr  "Snapshot USA 2019" "Snapshot USA 2019" "Snapshot USA 2019" "Snapshot USA 2019" ...
##  $ Camera_Trap_Array: chr  "Abilene" "Abilene" "Abilene" "Abilene" ...
##  $ Site_Name        : chr  "TX_Grassland_Abilene_10" "TX_Grassland_Abilene_11" "TX_Grassland_Abilene_12" "TX_Grassland_Abilene_2" ...
##  $ Deployment_ID    : chr  "TX_Grassland_Abilene_10" "TX_Grassland_Abilene_11" "TX_Grassland_Abilene_12" "TX_Grassland_Abilene_2" ...
##  $ Start_Date       : chr  "9/3/2019" "9/4/2019" "9/1/2019" "9/1/2019" ...
##  $ End_Date         : chr  "11/2/2019" "11/2/2019" "10/12/2019" "11/2/2019" ...
##  $ Survey_Nights    : int  60 59 41 62 61 57 53 60 39 21 ...
##  $ Latitude         : num  32.2 32.2 32.2 32.2 32.2 ...
##  $ Longitude        : num  -99.9 -99.9 -99.9 -99.9 -99.9 ...
##  $ Habitat          : chr  "Grassland" "Grassland" "Grassland" "Grassland" ...
##  $ Development_Level: chr  "Rural" "Rural" "Rural" "Rural" ...
##  $ Feature_Type     : chr  "None" "None" "None" "None" ...
```

```
str(recs_all)
```

```
## 'data.frame':    972415 obs. of  16 variables:
##  $ Year             : int  2019 2019 2019 2019 2019 2019 2019 2019 2019 2019 ...
##  $ Project          : chr  "Snapshot USA 2019" "Snapshot USA 2019" "Snapshot USA 2019" "Snapshot USA 2019" ...
##  $ Camera_Trap_Array: chr  "VermilionCliffs" "VermilionCliffs" "VermilionCliffs" "VermilionCliffs" ...
##  $ Deployment_ID    : chr  "AZ_Grassland_Vermilion_Cliffs_11" "AZ_Grassland_Vermilion_Cliffs_11" "AZ_Grassland_Vermilion_Cliffs_11" "AZ_Grassland_Vermilion_Cliffs_11" ...
##  $ Sequence_ID      : chr  "d63190s2" "d63190s5" "d63190s10" "d63190s11" ...
##  $ Start_Time       : chr  "8/25/2019 8:32:00" "8/29/2019 8:02:00" "9/1/2019 7:23:00" "9/2/2019 7:49:00" ...
##  $ End_Time         : chr  "8/25/2019 8:32:00" "8/29/2019 8:02:00" "9/1/2019 7:23:00" "9/2/2019 7:49:00" ...
##  $ Class            : chr  "Mammalia" "Mammalia" "Mammalia" "Mammalia" ...
##  $ Order            : chr  "Rodentia" "Rodentia" "Rodentia" "Rodentia" ...
##  $ Family           : chr  "Sciuridae" "Sciuridae" "Sciuridae" "Sciuridae" ...
##  $ Genus            : chr  "Sciurus" "Sciurus" "Sciurus" "Sciurus" ...
##  $ Species          : chr  "aberti" "aberti" "aberti" "aberti" ...
##  $ Common_Name      : chr  "Abert's Squirrel" "Abert's Squirrel" "Abert's Squirrel" "Abert's Squirrel" ...
##  $ Age              : chr  "Unknown" "Unknown" "Unknown" "Unknown" ...
##  $ Sex              : chr  "Unknown" "Unknown" "Unknown" "Unknown" ...
##  $ Group_Size       : int  1 1 1 1 1 1 1 1 1 1 ...
```

Next we subset the data to the survey called “Hebblewhite” in year
2022, a camera trapping survey conducted at 31 locations in Lubrecht
Experimental Forest, Montana, between September and December of
2022.

```
trap_array <- "Hebblewhite"
survey_year <- 2022

cams <- cams_all[cams_all$Camera_Trap_Array == trap_array & cams_all$Year == survey_year,]
recs <- recs_all[recs_all$Camera_Trap_Array == trap_array & recs_all$Year == survey_year,]
```

Camera trap stations can also be filtered in the dashboard in the
“Filters” tab, but this is done here via code for clarity and to for
performance.

In this particular dataset, we add the `Site_Name` column
to the record table as a shared station identifier between the camera
and record table (deployment ID will be used like a camera ID because
sometimes there are several deployments per site).

```
recs <- merge(recs, cams[, c("Deployment_ID", "Site_Name")])
```

### Dashboard-based workflow

The most basic approach for loading a dataset in the dashboard is to
open the empty dashboard and use the Import dialog for loading the
CSVs.

```
surveyDashboard()
```

The downside is that the user needs to provide the specification of
the tables in the UI, which can be tedious, especially when done
multiple times. To streamline data loading it is advisable to specify
the particulars of the data set in the call to
`surveyDashboard` instead. In particular, we provide the
input tables, column names of the tables, date/time formats and the
coordinate system.

```
surveyDashboard(
  # input tables
  CTtable = cams, 
  recordTable = recs, 
  
  # column names
  stationCol = "Site_Name",
  cameraCol = "Deployment_ID",
  setupCol = "Start_Date",
  retrievalCol = "End_Date",
  recordDateTimeCol = "Start_Time",
  speciesCol = "Common_Name",
  xcol = "Longitude",
  ycol = "Latitude",
  
  # treat cameras as independent?
  camerasIndependent = TRUE,
  
  # coordinate system
  crs = 4326,  

  # date / time formats
  CTdateFormat = "mdy",
  recordDateTimeFormat = "mdy HMS"
  )
```

In the dashboard, the user can explore the data, filter as needed,
visualize species activity patterns, add covariates, run and analyse
single and multi-species occupancy models, and generate and visualize
predictions from these. Using the dashboard does not require any R
code.

```
knitr::include_graphics(path = "figures/dashboard_screenshot1.png")
```

Screenshot of surveyDashboard()’s summary page.

### Code-based workflow

Alternatively, users can perform all analyses using traditional R
code. Below we outline the workflow for some common analyses using the
example data set.

#### Defining data set properties

Defining the data set properties upfront makes it easier to adapt the
code below to other data sets.

```
# set the column names of dataset

# in both station and record table: 
stationCol <- "Site_Name"
cameraCol  <- "Deployment_ID"

# in camera table
setupCol     <- "Start_Date"
retrievalCol <- "End_Date"
xcol         <- "Longitude"
ycol         <- "Latitude"

# in record table
speciesCol           <- "Common_Name"
recordDateTimeCol    <- "Start_Time"
recordDateTimeFormat <- "mdy HMS"
dateFormat           <- "mdy"


# set some other information about dataset
timeZone <- "America/New_York"  # Set appropriate time zone for data set
crs      <- "EPSG:4326"   # coordinate system of xcol/ycol: WGS84 (latlong)
```

By making `cams` a spatial object we can plot it easily in
an interactive map.

```
cams_sf <- st_as_sf(cams, 
                    coords = c(xcol, ycol), 
                    crs = crs)  

mapview(cams_sf)
```

#### Temporal independence of records

Camera trap records can be filtered for temporal independence. Sixty
minutes are commonly used. While filtering for temporal independence
generally does not affect occupancy estimates, it is good practice for
activity-based analyses.

```
recs <- filterRecordTable(recordTable = recs,
                  minDeltaTime = 60, 
                  deltaTimeComparedTo = "lastIndependentRecord",
                  speciesCol = speciesCol,
                  stationCol = stationCol,
                  cameraCol = cameraCol,
                  camerasIndependent = T,
                  recordDateTimeCol = recordDateTimeCol,
                  recordDateTimeFormat = recordDateTimeFormat,
                  quiet = TRUE)
```

```
## timeZone is not specified. Assuming UTC
```

Note that `filterRecordTable` converts the date/time
column in the record table to the standard format (YYYY-MM-DD HH:MM:SS),
hence it is not necessary to specify `recordDateTimeFormat`
below.

#### Adding covariates to data set

We can easily add site covariates to the data set for use in
ecological analyses below. In this example we download a digital
elevation model and derive some terrain metrics (topographic position,
ruggedness, slope). This requires an internet connection.

Before creating covariates though it makes sense to aggregate the
camera trap table to station level (currently it is station-deployment).
This can also be done after extracting covariates, but conceptually it
makes more sense beforehand.

```
cams_sf_agg <- aggregateStations(CTtable = cams_sf, 
                   stationCol = stationCol,
                   cameraCol = cameraCol,
                   setupCol = setupCol,
                   retrievalCol = retrievalCol,
                   dateFormat = dateFormat)
```

Now we extract the covariates at the camera trap locations (requires
internet connection when `download_elevation = TRUE`).

```
covariates <- createCovariates(CTtable = cams_sf_agg,
                               # directory = dir_data,  # local raster, in same folder as tables after unzipping
                               download_elevation = TRUE,    # online DEM
                               terrain_measures = c("TPI", "TRI", "slope"),
                               buffer_ct = 100,    # extract covariate values in buffer around cameras (meters)
                               resolution = 200,   # meter (prediction raster pixel size))
                               standardize_na = TRUE,
                               scale_covariates = TRUE)   # provides scaled covariates
```

The function returns the camera trap table with added covariates on
the original scale and (because `scale_covariates = TRUE`)
standardized to mean = 0 and standard deviation = 1.

```
# extract the camera trap table with the original covariates added
covs_model_orig <- covariates$CTtable

# extract the camera trap table with the new (scaled) covariates added
covs_model_scaled <- covariates$CTtable_scaled
```

The `createCovariates` function provides not only the
camera trap table with covariates added, but also rasters for use in
predictions.

Plot the extracted covariate raster for predictions (with camera trap
locations overlaid).

```
prediction_stack <- covariates$predictionRaster
plot(prediction_stack, fun = function() points(vect(cams_sf_agg)))
```

Original (unscaled) covariate rasters

```
prediction_stack_scaled <- covariates$predictionRaster_scaled
plot(prediction_stack_scaled, fun = function() points(vect(cams_sf_agg)))
```

Original (unscaled) covariate rasters (mean = 0, SD = 1)

Alternative ways for adding covariates to the camera trap table
include:

- providing custom rasters to `createCovariates` which are
  then processed in a standardized way
- adding covariate columns to the camera trap table manually (only
  allows for predictions at camera trap sites).

#### Creating a camera operation matrix

The camera operation matrix shows the daily operational status of
each camera trap. It is needed for accurately calculating sampling
effort for occupancy models.

```
camop <- cameraOperation(
  CTtable = cams,
  stationCol = stationCol,
  cameraCol = cameraCol,
  setupCol = setupCol,
  retrievalCol = retrievalCol,
  dateFormat = dateFormat,
  byCamera = FALSE,  # Aggregate data from multiple cameras at same station
  allCamsOn = FALSE, # Don't require all cameras to be functioning
  camerasIndependent = TRUE # Important: Treat cameras at same station as independent? *
)
```

```
# plot camera operation matrix
camtrapR:::camopPlot(camop)
```

Camera operation matrix

#### Basic ecological analyses

##### Species activity

One can compute single-species activity density using the
`overlap` package, as shown here for pumas.

```
activityDensity(recordTable = recs,
                species = "Puma",
                speciesCol = speciesCol,
                recordDateTimeCol = recordDateTimeCol)
```

Single-species activity density of Puma.

One can also compute two-species activity overlaps.

```
activityOverlap(recordTable = recs,
                speciesA = "Puma",
                speciesB = "American Black Bear",
                speciesCol = speciesCol,
                recordDateTimeCol = recordDateTimeCol)
```

Two-species activity overlap between Puma and American Black Bear.

##### Species accumulation curves

Various types of species accumulation curves can be created by
leveraging and augmenting the functionalities of the `iNEXT`
package.

The standard approach implemented in `iNEXT` is to plot
species diversity as a function of the number of samling locations,
hence analyzing the spatial aspect of species accumulation. It is
implemented by setting `x_unit = "station"`.

```
specaccum1 <- speciesAccum(CTtable = cams_sf_agg,
             recordTable = recs,
             speciesCol = speciesCol,
             recordDateTimeCol = recordDateTimeCol,
             setupCol = setupCol,
             stationCol = stationCol,
             x_unit = "station")

plot(specaccum1)
```

camtrapR furthermore offers a temporal analyses of species diversity
by plotting diversity as a function of survey duration.

The analysis can be conducted by station (beginning with the first
survey day *at each station separately*, by setting `x\_unit =
“station\_day”).

```
specaccum2 <- speciesAccum(CTtable = cams_sf_agg,
             recordTable = recs,
             speciesCol = speciesCol,
             recordDateTimeCol = recordDateTimeCol,
             dateFormat = "ymd",
             setupCol = setupCol,
             stationCol = stationCol,
             x_unit = "station_day")

plot(specaccum2)
```

Alternatively, the analysis can be conducted for the entire survey
duration (beginning with the first day of the survey, by setting `x\_unit
= “survey\_day”).

```
specaccum3 <- speciesAccum(CTtable = cams_sf_agg,
             recordTable = recs,
             speciesCol = speciesCol,
             recordDateTimeCol = recordDateTimeCol,
             dateFormat = "ymd",
             setupCol = setupCol,
             stationCol = stationCol,
             x_unit = "survey_day")

plot(specaccum3)
```

##### Single-species occupancy models

Single-species occupancy models can be fit with (for example)
packages `unmarked` or `ubms`.

First we create a detection history of the target species, here, the
red fox.

```
dethist_fox <- detectionHistory(recordTable = recs,
                 species = "Red Fox",
                 camOp = camop,
                 stationCol = stationCol,
                 speciesCol = speciesCol,
                 recordDateTimeCol = recordDateTimeCol,
                 occasionLength = 10)
```

```
## Warning: timeZone is not specified. Assuming UTC
```

Then we combine the detection history, site covariates and
observation covariates in an `unmarkedFrameOccu`.

```
umf_fox <- unmarkedFrameOccu(y = dethist_fox$detection_history,
                  siteCovs = st_drop_geometry(covs_model_orig),
                  obsCovs = list(effort = dethist_fox$effort))
```

```
## Warning: siteCovs contains characters. Converting them to factors.
```

From here on users can simply follow the standard
`unmarked` or `ubms` workflows.

For example, one could fit the following models\_

```
occu0 <- occu(~effort ~1, data = umf_fox)
occu1 <- occu(~effort ~ scale(TPI), data = umf_fox)
```

And look at the parameter estimates.

```
occu0
```

```
## 
## Call:
## occu(formula = ~effort ~ 1, data = umf_fox)
## 
## Occupancy (logit-scale):
##  Estimate    SE    z P(>|z|)
##      0.58 0.398 1.46   0.145
## 
## Detection (logit-scale):
##             Estimate    SE     z P(>|z|)
## (Intercept)   -2.999 1.033 -2.90 0.00369
## effort         0.246 0.107  2.29 0.02189
## 
## AIC: 265.3803 
## Number of sites: 31
```

```
occu1
```

```
## 
## Call:
## occu(formula = ~effort ~ scale(TPI), data = umf_fox)
## 
## Occupancy (logit-scale):
##             Estimate    SE     z P(>|z|)
## (Intercept)    0.611 0.424  1.44    0.15
## scale(TPI)    -0.748 0.481 -1.55    0.12
## 
## Detection (logit-scale):
##             Estimate    SE     z P(>|z|)
## (Intercept)   -2.983 1.032 -2.89 0.00386
## effort         0.244 0.107  2.28 0.02263
## 
## AIC: 264.42 
## Number of sites: 31
```

And predict occupancy probabilities across the landscape (only for
illustration).

```
pred_occu_fox <- predict(occu1, 
        type = "state",
        newdata = prediction_stack)
```

```
plot(pred_occu_fox$Predicted)
```

Predicted occupancy probability of Red Fox

#### Multi-species occupancy models

Multi-species (community) occupancy models in `camtrapR`
are currently available in a Bayesian framework using either JAGS or
NIMBLE. Future versions may also offer support for the frequentist
implementation in `unmarked`.

##### Species selection

In this demonstration we select species with at least 3
detections.

```
min_n_detections <- 3
```

Furthermore we remove human, vehicle, domestic animal, and
unidentified animal detections.

```
species_counts <- table(recs[, speciesCol])

# remove undesired records 
species_to_ignore <- paste(c("Human", "Domestic", "Species", "Family", "Vehicle", "Unknown", "Mammal", "Bird", "Animal"), collapse = "|")
species_shortlist <- stringr::str_subset(names(species_counts), pattern = species_to_ignore, negate = TRUE)

# subset to shortlisted species
species_counts <- species_counts[names(species_counts) %in% species_shortlist]

# remove species with too few records
species_counts <- species_counts[species_counts >= min_n_detections]

target_species <- names(species_counts)
```

That leaves 19 species for the community occupancy model:

Note, filtering species is easier on the dashboard.

##### Preparing occupancy model input

Next, we create a list of detection histories for all target
species.

```
DetHist_list <- detectionHistory(
  recordTable = recs,
  camOp = camop,
  stationCol = stationCol,
  speciesCol = speciesCol,
  recordDateTimeCol = recordDateTimeCol,
  species = target_species,     # multiple species at once
  occasionLength = 10,          # 10-day occasions
  day1 = "station",
  datesAsOccasionNames = FALSE,
  includeEffort = TRUE,
  scaleEffort = FALSE,
  timeZone = timeZone
)
```

Finally, we bundle the data for the `communityModel`
function. l. We combine the detection histories, covariate data frame
and observation covariate effort (returned by
`detectionHistory`).

```
data_list <- list(
  ylist = DetHist_list$detection_history,
  siteCovs = covs_model_scaled,
  obsCovs = list(effort = DetHist_list$effort)  # effort is identical for all species
)
```

##### Creating and fitting community occupancy models

Users can flexibly define their desired model structure and the
`communityModel` function writes JAGS/NIMBLE code for it. It
also automatically chooses start values for the MCMC simulation and sets
parameters to monitor.

In this example we create a community occupancy model including
a:

- fixed effect of effort on detection probability
- species-specific random effects of elevation, ruggedness (TRI) and
  topographic position index (TPI) on occupancy probability.

For more possibilities (like nested random effects, or
station-species specific random effects for detection probability)
please see the vignette about Multi-species occupancy models.

```
modelfile1 <- file.path(getwd(), "community_model1.txt")

mod1 <- communityModel(
  data_list,
  occuCovs = list(ranef = c("elevation", "TRI", "TPI")),
  detCovsObservation = list(fixed = "effort"),
  intercepts = list(det = "ranef", occu = "ranef"),
  modelFile = modelfile1
)
```

```
## Wrote model to C:/Arbeit/camtrapR/camtrapR_supplement/community_model1.txt
```

Now we are ready to fit the model. The In the `fit()`
function users can specify the MCMC sampling: number of samples,
burn-in, number of chains, thinning.

```
# number of posterior draws
# this is a low number for demonstration
n_draws_fit <- 2000  

# Fit the model
fit1 <- fit(
  mod1,
  n.iter = n_draws_fit
)
```

```
## Compiling model graph
##    Resolving undeclared variables
##    Allocating nodes
## Graph information:
##    Observed stochastic nodes: 5168
##    Unobserved stochastic nodes: 2006
##    Total graph size: 15471
## 
## Initializing model
## 
## NOTE: Stopping adaptation
```

##### Model diagnostics

This is to ensure the chains converged and provide stable estimates;
diagnostics should be checked before visualizing results / making
predictions.

Note: this does not mean the models are good (fit the data well),
just that the chains converged to stable parameter estimates. The
dashboard contains interactive visualizations of the model fit.

###### Traceplots

Traceplots are important diagnostic tools showing chain mixing and
distribution of parameter estimates.

The easiest way to plot them is with `plot` from the coda
package, which is the default when plotting the model output. It does
not work well in markdown mode and is therefore not run here

```
# coda::plot() works well in interactive mode but not in markdown
plot(fit1)
```

Alternatively we can use the `bayesplot` package for
plotting traceplots (and other visualizations).

For example, here are the traceplots of the mean (community)
parameter estimates.

```
bayesplot::color_scheme_set(c("mix-blue-red"))
bayesplot::mcmc_trace(fit1, regex_pars = "mean")
```

Traceplots created with the bayesplot package.

###### Convergence

Gelman-Rubin convergence statistic (potential scale reduction factor)
estimates close to 1 indicate convergence; values > 1.1 indicate
non-convergence / incomplete mixing. The table contains model parameter
estimates (alpha, beta) and derived metrics (Nspecies).

```
gd1 <- coda::gelman.diag(fit1,  multivariate = FALSE)
```

###### Goodness of fit test

We can conduct a post-hoc goodness-of-fit test using posterior
predictive checks. First, we predict detection probability p and
occupancy probability p for 100 posterior samples. We set
`seed` to ensure the same samples are used in both
predictions.

```
# use same seed in both predict calls to ensure they use the same posterior samples
seed <- 100  

# run goodness-of-fit test for 100 posterior samples (low number for demonstration)
draws_gof <- 100
```

```
# Get predictions for p and psi using subset of MCMC draws
p_pred <- predict(
  object = mod1,
  mcmc.list = fit1,
  type = "p_array",
  draws = draws_gof,
  seed = seed
)

psi_pred <- predict(
  object = mod1,
  mcmc.list = fit1,
  type = "psi_array",
  draws = draws_gof,
  seed = seed
)
```

With both `p` and `psi` we can run the
community-level posterior predictive checks. The checks use model
residuals, here (specified by `type = "FT"`) Freeman-Tukey
residuals. Goodness of fit can be tested in various ways. Please see the
function help file and the instructions page for the Goodness-of-fit
test in the dashboard.

```
gof_results <- PPC.community(
  p = p_pred,
  psi = psi_pred,
  y = mod1@input$ylist,
  model = "Occupancy",
  type = "FT"
)
```

The function returns a table with Bayesian p-values (both by species
and for the entire community / the whole model).

```
bayesian_p_table <- gof_results$BP
rownames(bayesian_p_table) <- NULL
bayesian_p_overall <- bayesian_p_table$BP[bayesian_p_table$Species == "Community"]
```

Overall Bayesian p-value is 0.98, indicating lack of fit.

Species-level Bayesian p-values are:

```
DT::datatable(bayesian_p_table,
               caption = paste0("Table ", table_number, ": Bayesian p-values of community occupancy model, by species."))
```

These results indicate that the model does not adequately describe
the data, both overall and also for several individual species.
Specifically, since we set `z.cond = TRUE`, the detection
submodel shows lack of fit.As this is merely a demonstration, we will
proceed with the model as is. In a proper ecological analysis this would
be an opportunity to investigate the details of lack of fit (common
traits of the species with lack of fit, such as common vs. rare species,
or behavioural aspects) in order to improve the structure of the
detection submodel. Potential improvements are:

- using random slopes for the effect of effort, or
- adding more detection covariates (e.g. environmental,
  camera-related, or relate to human activity)

Both approaches - allowing more variation between species or
providing more explanatory variables - can help improve model fit.

##### Model parameter statistics

Posterior summaries of the parameter estimates are created with
`summary`. The following tables again contain both model
parameter estimates and derived metrics (like Nspecies).

```
fit1_summary <- summary(fit1)
```

```
DT::datatable(round(fit1_summary$statistics, 2),
              caption = paste0("Table ", table_number, ": Summary statistics of parameter estimates from multi-species occupancy model."))
```

```
DT::datatable(round(fit1_summary$quantiles, 2),
              caption = paste0("Table ", table_number, ": Quantiles of parameter estimates from multi-species occupancy model."))
```

##### Visualizing covariate effects

###### Marginal effect plots

Marginal effect plots (“reponse curves”) can be plotted with
`plot_effects`, both for the occupancy and the detection
submodel. They show how the covariates affect the response variable.

```
# Plot marginal effects for occupancy
camtrapR::plot_effects(
  object = mod1,
  mcmc.list = fit1,
  submodel = "state"
)
```

```
## $elevation
```

```
## 
## $TRI
```

```
## 
## $TPI
```

```
# Plot marginal effects for detection
camtrapR::plot_effects(
  object = mod1,
  mcmc.list = fit1,
  submodel = "det"
)
```

```
## $effort
```

###### Effect size plots

Effect size plots show the magnitude of coefficients and are suitable
for comparisons between species, or to see species effects relative to
community effects. They can be plotted with `plot_coef`, and
again are available for the occupancy and detection submodels.

```
# Plot effect sizes for occupancy
plot_coef(
  object = mod1,
  mcmc.list = fit1,
  submodel = "state"
)
```

```
## $elevation
```

```
## 
## $TRI
```

```
## 
## $TPI
```

```
# Plot effect sizes for detection
plot_coef(
  object = mod1,
  mcmc.list = fit1,
  submodel = "det"
)
```

```
## $effort
```

##### Spatial predictions

Spatial predictions can be created from the model object, model
output and the previously created prediction raster.

Since we used scaled covariates in the model we also need to use
scaled prediction rasters to match the scaled covariates used in the
model. These were provided automatically by
`createCovariates`.

Predictions can be memory-intensive and potentially time-consuming,
especially when predicting over large areas, using high resolution
rasters, and many posterior samples. While the function attempts to
prevent out-of-memory errors, we recommend starting with lower
resolution / fewer samples and gradually increase both.

###### Species occupancy probability

Species occupancy probability can be predicted by specifying
`type = "psi"`.

```
n_draws_predict <- 500   # this is a slightly low number to speed up computation and reduce memory load.
```

```
# Make spatial predictions for each species
predictions_psi <- predict(
  object = mod1,
  mcmc.list = fit1,
  x = prediction_stack_scaled,
  type = "psi",
  draws = n_draws_predict   
)
```

```
# Plot species occupancy maps
plot(predictions_psi$mean, 
     zlim = c(0, 1),
     col = hcl.colors(100),
     maxnl = 9)  # Show first 9 species only
```

Maps of predicted species occupanc probability (for the first nine
species)

```
# # show remaining species
# plot(predictions_psi$mean[[10:20]], 
#      zlim = c(0, 1),
#      col = hcl.colors(100))
```

###### Species richness

Species richness can be predicted by specifying
`type = "richness"`.

```
predictions_rich <- predict(
  object = mod1,
  mcmc.list = fit1,
  x = prediction_stack_scaled,
  type = "richness",
  draws = n_draws_predict
)
```

We can create a simple static map of mean predicted species richness
and its associated standard deviation.

```
plot(predictions_rich, 
     col = hcl.colors(100))
```

Maps of predicted species richness (mean prediction and standard
deviation).

And we can also plot the richness prediction on an interactive
map.

```
mapview(predictions_rich$mean,
        col.regions = hcl.colors(10),
        alpha.regions = 1) # no transparency
```

###### Percentage of area occupied

Percentage of area occupied can be predicted by specifying
`type = "pao"`.

```
predictions_pao <- predict(
  object = mod1,
  mcmc.list = fit1,
  x = prediction_stack_scaled,
  type = "pao",
  draws = n_draws_predict
)
```

```
## No id variables; using all as measure variables
## No id variables; using all as measure variables
```

```
DT::datatable(round(predictions_pao$pao_summary, 2),
              caption = paste0("Table ", table_number, ": Statistics of percentage of area occupied (PAO) by species."))
```

Violin plot of all species’predicted percentage of area occupied (PAO)
across the prediction raster

### Session information

```
sessionInfo()
```

```
## R version 4.5.1 (2025-06-13 ucrt)
## Platform: x86_64-w64-mingw32/x64
## Running under: Windows 10 x64 (build 19045)
## 
## Matrix products: internal
##   LAPACK version 3.12.1
## 
## locale:
## [1] LC_COLLATE=German_Germany.utf8  LC_CTYPE=German_Germany.utf8   
## [3] LC_MONETARY=German_Germany.utf8 LC_NUMERIC=C                   
## [5] LC_TIME=German_Germany.utf8    
## 
## time zone: UTC
## tzcode source: internal
## 
## attached base packages:
## [1] stats     graphics  grDevices utils     datasets  methods   base     
## 
## other attached packages:
## [1] DT_0.33          bayesplot_1.13.0 ggplot2_3.5.2    readr_2.1.5     
## [5] mapview_2.11.2   unmarked_1.5.0   terra_1.8-60     sf_1.0-21       
## [9] camtrapR_2.6.0  
## 
## loaded via a namespace (and not attached):
##   [1] RColorBrewer_1.1-3      tensorA_0.36.2.1        rstudioapi_0.17.1      
##   [4] jsonlite_2.0.0          wk_0.9.4                magrittr_2.0.3         
##   [7] farver_2.1.2            iNEXT_3.0.2             rmarkdown_2.29         
##  [10] vctrs_0.6.5             base64enc_0.1-3         RcppNumerical_0.6-0    
##  [13] htmltools_0.5.8.1       progress_1.2.3          distributional_0.5.0   
##  [16] curl_7.0.0              rjags_4-17              raster_3.6-32          
##  [19] s2_1.1.9                Formula_1.2-5           sass_0.4.10            
##  [22] slippymath_0.3.1        KernSmooth_2.23-26      bslib_0.9.0            
##  [25] htmlwidgets_1.6.4       plyr_1.8.9              lubridate_1.9.4        
##  [28] stars_0.6-8             cachem_1.1.0            uuid_1.2-1             
##  [31] mime_0.13               lifecycle_1.0.4         pkgconfig_2.0.3        
##  [34] elevatr_0.99.0          Matrix_1.7-3            R6_2.6.1               
##  [37] fastmap_1.2.0           rbibutils_2.3           shiny_1.11.1           
##  [40] digest_0.6.37           colorspace_2.1-1        leafem_0.2.4           
##  [43] textshaping_1.0.1       crosstalk_1.2.1         Hmisc_5.2-3            
##  [46] suntools_1.0.1          labeling_0.4.3          progressr_0.15.1       
##  [49] timechange_0.3.0        httr_1.4.7              abind_1.4-8            
##  [52] mgcv_1.9-3              compiler_4.5.1          proxy_0.4-27           
##  [55] withr_3.0.2             htmlTable_2.4.3         brew_1.0-10            
##  [58] backports_1.5.0         DBI_1.2.3               MASS_7.3-65            
##  [61] leaflet_2.2.2           classInt_0.4-11         tools_4.5.1            
##  [64] units_0.8-7             foreign_0.8-90          httpuv_1.6.16          
##  [67] nnet_7.3-20             glue_1.8.0              satellite_1.0.6        
##  [70] nlme_3.1-168            promises_1.3.3          grid_4.5.1             
##  [73] checkmate_2.3.3         cluster_2.1.8.1         reshape2_1.4.4         
##  [76] generics_0.1.4          gtable_0.3.6            leaflet.providers_2.0.0
##  [79] tzdb_0.5.0              shinyBS_0.61.1          class_7.3-23           
##  [82] data.table_1.17.8       hms_1.1.3               overlap_0.3.9          
##  [85] sp_2.2-0                pillar_1.11.0           stringr_1.5.1          
##  [88] posterior_1.6.1         later_1.4.3             splines_4.5.1          
##  [91] dplyr_1.1.4             lattice_0.22-7          tidyselect_1.2.1       
##  [94] pbapply_1.7-4           knitr_1.50              gridExtra_2.3          
##  [97] reformulas_0.4.1        svglite_2.2.1           stats4_4.5.1           
## [100] xfun_0.52               shinydashboard_0.7.3    leafpop_0.1.0          
## [103] stringi_1.8.7           yaml_2.3.10             evaluate_1.0.4         
## [106] codetools_0.2-20        tibble_3.3.0            cli_3.6.5              
## [109] rpart_4.1.24            RcppParallel_5.1.10     xtable_1.8-4           
## [112] systemfonts_1.2.3       Rdpack_2.6.4            jquerylib_0.1.4        
## [115] secr_5.2.4              Rcpp_1.1.0              coda_0.19-4.1          
## [118] png_0.1-8               parallel_4.5.1          prettyunits_1.2.0      
## [121] mvtnorm_1.3-3           scales_1.4.0            e1071_1.7-16           
## [124] purrr_1.1.0             crayon_1.5.3            rlang_1.1.6
```
